# Dynamic knowledge representation of blood brain barrier activation, injury and restitution with a novel agent-based model

**DOI:** 10.64898/2026.07.20.739661

**Authors:** Gary An, Chase Cockrell

## Abstract

The blood brain barrier (BBB) tightly regulates the interface between the central nervous system and the systemic circulation. The function of the BBB is the output of an architecturally complex, multicellular neurovascular unit that consists of brain microvascular endothelial cells, pericytes, astrocytes, microglia and parenchymal neurons. Dysfunction of the BBB has been linked to numerous neurological diseases, such as multiple sclerosis, Alzheimer’s Disease, traumatic brain injury/chronic traumatic encephalopathy and stroke. Also, the control of the BBB over permeability makes it an ongoing interest and target for pharmaceutical development. Herein we present Blood Brain Barrier Agent-based Model (BBBABM), the first computational model that mechanistically represents the cellular components of the neurovascular unit and the molecular interactions that govern the response of the BBB to injury/activation, including the restorative functions that provide baseline homeostasis regarding the health of the BBB. Simulation experiments with the BBBABM replicate expected dynamics of disruption and restoration that demonstrate a dose responsiveness to the severity of the initial insult. The development of the BBBABM provided insight into the contribution of various forms of cellular-molecular responses to overall BBB dysfunction, including integrating across time scales from minutes to weeks. This initial implementation of the BBBABM offers numerous future paths for development, including being able to mechanistically represent the long time scales (years/decades) present in the pathophysiology of chronic neurological diseases and providing a platform to enhance therapeutic development via in silico trials and as the basis for mechanistic cellular-molecular Digital Twins.

## Introduction

The blood brain barrier (BBB) represents a unique multi-cellular unit (sometimes termed the neurovascular unit (Nguyen, Pham et al. 2026)), that controls and mediates the communication between the systemic circulation and the central nervous system. Composed of specialized endothelial cells, pericytes, astrocytes and microglial cells, the BBB is of high interest both in terms of pathophysiology and pharmacology. Dysfunction of the BBB has been invoked in a host of disease processes, ranging across infectious disease (Kim 2008, Galea 2021), multiple sclerosis (Nishihara, Perriot et al. 2022, Zierfuss, Larochelle et al. 2024), Alzheimer’s Disease (Montagne, Zhao et al. 2017, Sweeney, Sagare et al. 2018), stroke (Abdullahi, Tripathi et al. 2018, Yang, Hawkins et al. 2019) and traumatic brain injury (TBI) (Shlosberg, Benifla et al. 2010, Chodobski, Zink et al. 2011, Hay, Johnson et al. 2015, Price, Wilson et al. 2016, Cash and Theus 2020, Sulhan, Lyon et al. 2020, van Vliet, Ndode-Ekane et al. 2020), including chronic traumatic encephalopathy (CTE) (Doherty, O’Keefe et al. 2016, Greene, Brennan et al. 2026). Many of these diseases involve some degree of neuroinflammation that is initiated and/or propagated by activation of the endothelial component of the BBB to a point where baseline homeostatic processes take on pathological trajectories. Both the normal compensatory and pathophysiological responses are dynamic processes, and characterizing the shift from a healthy state to one of disease requires the ability to represent these dynamics as they emerge from the mechanistic understanding generated by experimental investigations. We term this process as “dynamic knowledge representation” and have extensively utilized agent-based models (ABMs) for dynamic knowledge representation such that the behavioral consequences of a particular mechanism-based hypothesis structure can be visualized, interrogated and evaluated (An 2008, An 2009, An 2015). Herein we present what is, to our knowledge, the first mathematical/computational model, the Blood Brain Barrier Agent-based Model (BBBABM), that mechanistically represents the activation, injury and restoration of the BBB (for survey of existing methods see (Jafarabadi, Datta et al. 2026)).

## 1. Methods

### 1.1 Development of BBBABM

The BBBABM is an ABM that includes the cellular components of the BBB, the functions associated with the BBB’s response to Brain Microvascular Endothelial Cell (BMEC) injury/activation, and the restorative functions that provide baseline homeostasis of the BBB. The process of developing the BBBABM follows our well-established workflow that involves a comprehensive literature review, with an emphasis on capturing robust and non-controversial features of the system of interest, the generation of a knowledge graph with an abstraction level suited for the modeling project and subsequent instantiation of that knowledge graph into an ABM (An 2008, An 2009, An 2015), in this case using the toolkit NetLogo(Wilensky 1999). The model can be found at https://github.com/Gary-An/Blood-Brain-Barrier-ABM.

### 1.2 Simulation Experiments

#### 1.2.1 Parameter Sweep of Trauma Severity

After creation of the BBBABM validation involve a series of simulation experiments to determine whether the model behaves in a plausible fashion (“face validity”). At the current level of development the goal is to demonstrate that the BBBABM behaves at this level. Validation simulations were performed via a parameter sweep of “Trauma Severity” value with 10 stochastic replicates per value. This is done to confirm dose-dependent responses of BBB components over a span of 14 days following insult. BBBABM behavior evaluated by trajectories of model outputs that representing 1) pathophysiologically-relevant system-level behavior: overall tight junction (TJ)-expression level and Permeability and 2) mediators chosen to reflect key implemented processes related to neuroinflammation: TNFa, MMP and leukocyte dynamics. Additional goal is to identify potential inflection points in terms of BBBABM behavior that will allow for categorization of degree of “Trauma Severity” that would correlate with minimal insult.

## 2. Results

### 2.1 Model Description

#### 2.1.1 Overview

The BBBABM is implemented in Netlogo and depicts a 2D cross-section of a neurovascular unit, representing the cellular components and spatial architecture that contribute to the BBB. The aggregate behaviors targeted by the BBBABM include: 1) baseline maintenance under healthy conditions, including effective recovery from endothelial activation below some threshold, 2) pathological behavior, in terms of permeability failure, tissue edema and neuroinflammation, when challenged with an insult greater than can be handled in the homeostatic condition and 3) the ability to recover from these more severe insults. The BBBABM reproduces several hallmark features of BBB injury and repair documented in the traumatic-brain-injury (TBI) and stroke literature: acute TJ collapse, cytotoxic and vasogenic edema, an inflammatory cascade with leukocyte recruitment, irreversible loss of endothelial cells at high injury severity, a characteristic two-phase (biphasic) increase in BBB permeability, and staged recovery driven by eight distinct biological mechanisms operating on timescales from minutes to weeks (Chodobski, Zink et al. 2011, Tietz and Engelhardt 2015, Bellut, Papp et al. 2021, van Hameren, Aboghazleh et al. 2024, Chagnot and Montagne 2025).

#### 2.1.2 Spatial-Temporal Properties

The time scale of the BBBABM is such that one iteration (“tick”) of the model represents 1 minute of real time. As noted above the BBBABM represents a 2D cross section of a neurovascular unit and does this with a 2D square grid that is 33 grid-spaces wide (x coordinates = -16 to 16) and 17 grid-spaces tall (y coordinates = 0 to 16), wrapping horizontally so the left and right edges are continuous. Moving from bottom to top, the patches are organized into biological zones:

- Blood lumen (vascular space), the bottom rows ycor < 5.
- The BMEC (Brain Microvascular Endothelial Cell) layer: a single row of 33 brain microvascular endothelial cells, one per column, at ycor = 5. This row is the barrier itself.
- Basement membrane / pericyte layer, just above the endothelium, ycor = 6.
- Astrocyte endfeet layer, ycor = 7.
- CNS parenchyma (neurons, microglia, perivascular macrophages, and leukocytes that have crossed the barrier), the upper rows ycor > 7.

This vertical arrangement follows the standard anatomy of the neurovascular unit, in which endothelial tight junctions, pericytes, and astrocyte endfeet jointly form the barrier (Shlosberg, Benifla et al. 2010, Tietz and Engelhardt 2015, Benarroch 2023, Nguyen, Pham et al. 2026). Because there is exactly one BMEC per column and cells are neither created nor destroyed at runtime, the barrier layer always contains 33 cells. Any BMEC that dies stays in place as a non-functional gap (more on this in Section 3.1.5.7 on Pyroptosis). Figure 1 is a schematic of the spatial organization of the BBBABM, its cellular components, included mediators and key recovery mechanisms to be described in the sections below.

**Figure 1:**
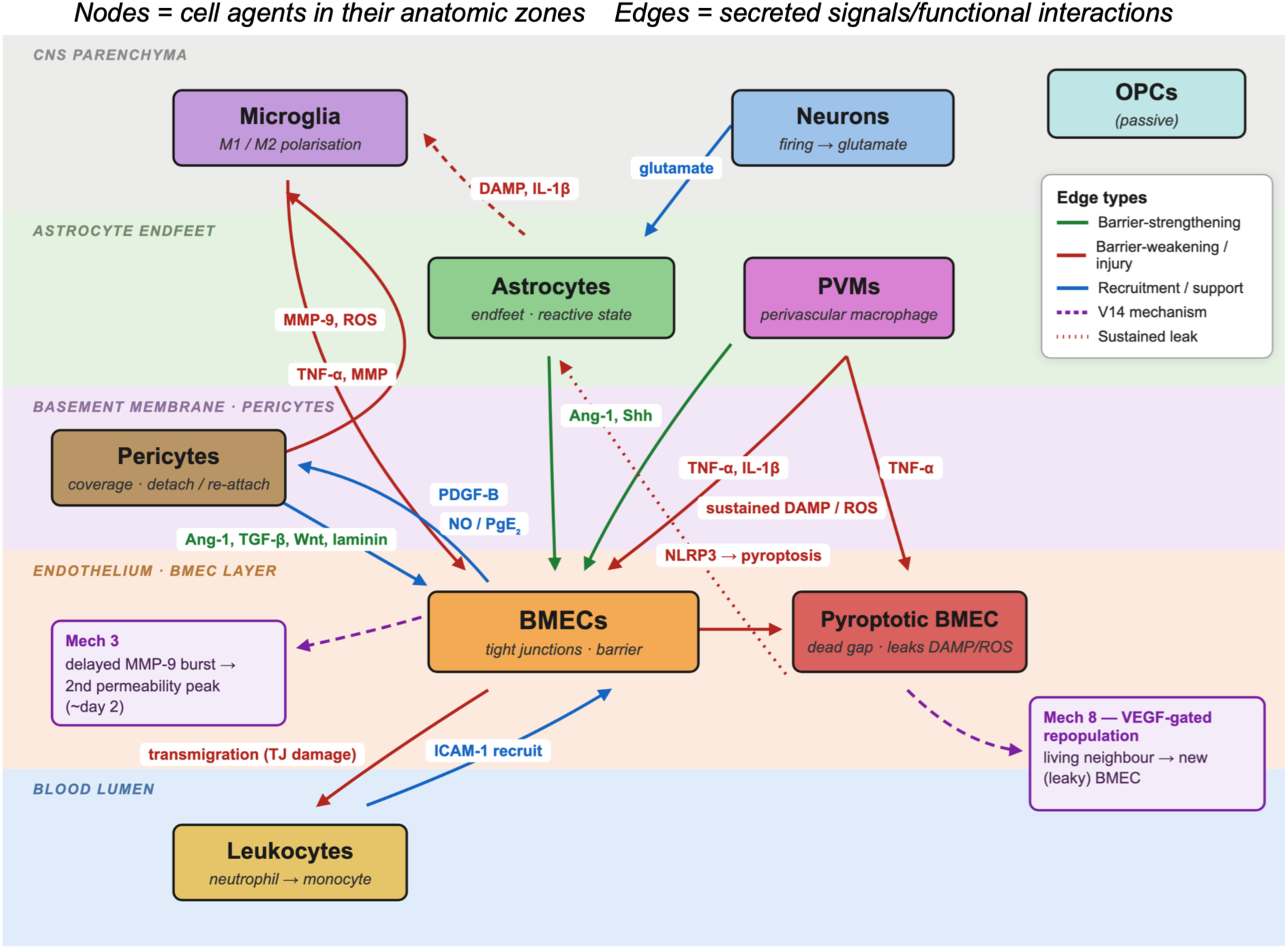
Schematic of Spatial Organization of the BBBABM.

#### 2.1.3 Cellular Agents

As an ABM the BBBABM represents specific cell-types as classes of computational agents. These agent-classes function as finite state machines that incorporate behavioral rules derived from the mechanistic information generated by basic science experimental biology; this is the information extracted from the literature and organized into a knowledge graph. It is via this property that ABMs can dynamically represent a given hypothesis structure regarding the behavior of a particular cell type. While the rules for a particular agent-class are shared by all individual agents created within that class, individual agent (cells) will have varying behavioral trajectories based on heterogeneity present in their local environment, mimicking how real cells respond and behave. The following cell types are represented in the BBBABM:

- BMECs: These represent the barrier function of the BBB. These agents have intrinsic polarity, where the basement membrane is at the “top” of each BMEC. Each tick, every living BMEC reads the chemical concentrations on its patch, updates its receptors, and recomputes its TJ expression from a balance of junction-strengthening signals (angiopoietin-1, sonic hedgehog, Wnt, and a claudin-5 synthesis term) against junction-weakening signals (TNF-*a*, matrix metalloproteinase, thrombin/PAR-1 signaling, IL-1beta, reactive oxygen species, and NF-kB activity) (Shlosberg, Benifla et al. 2010, Chodobski, Zink et al. 2011, Tietz and Engelhardt 2015, Bellut, Papp et al. 2021, Benarroch 2023, Nguyen, Pham et al. 2026). Each cell’s permeability is then derived from its TJ expression plus contributions from caveolar transcytosis, local VEGF, ROS damage, and local MMP. Cells also manage Weibel-Palade body exocytosis, ICAM-1 display for leukocyte capture, and an NLRP3 inflammasome that can trigger cell death (pyroptosis) (Birukova, Zagranichnaya et al. 2007, Liu, Li et al. 2018, Bellut, Papp et al. 2021).
- Pericytes: These agents wrap the vessel from the basement-membrane side (superior edge). They provide paracrine support (Ang-1, TGF-beta, Wnt), synthesize basement-membrane laminin, and contract or dilate in response to vasoactive signals. Under injury they can detach, and they re-attach and rebuild coverage slowly during recovery (Hori, Ohtsuki et al. 2004, Dohgu, Takata et al. 2005, Armulik, Genové et al. 2010, Chodobski, Zink et al. 2011).
- Astrocytes: These agents extend endfeet onto the vessel and couple neuronal activity to the barrier, secreting Ang-1, Shh, and nitric oxide. They become reactive under inflammation, shifting their output toward VEGF (Abbott, Rönnbäck et al. 2006, Chodobski, Zink et al. 2011).
- Neurons: Their primary function in the BBBABM fire at a tunable rate and release glutamate, driving neurovascular coupling (Chodobski, Zink et al. 2011, Galea 2021, Chagnot and Montagne 2025).
- Microglia. These mobile agents survey for inflammatory signals and damage-associated molecular patterns (DAMPs). Above a damage threshold they polarize to a pro-inflammatory M1 state and secrete TNF-*a* and MMP; below it they adopt a reparative M2 state (Chodobski, Zink et al. 2011, Galea 2021, Nguyen, Pham et al. 2026).
- Perivascular macrophages (PVMs): These mobile agents patrol the basement-membrane/endfeet interface and secrete TNF-*a* when activated (Chodobski, Zink et al. 2011, Galea 2021, Nguyen, Pham et al. 2026).
- Leukocytes: These mobile agents only appear in the BBBABM when they are recruited from the blood when the endothelium displays ICAM-1. This represents the process by which leukocytes are engaged and roll, adhere, and transmigrate across the barrier into the CNS parenchyma, where they release MMP-9 and ROS before dying (Chodobski, Zink et al. 2011, Galea 2021, Nguyen, Pham et al. 2026). They provide the visible cellular arm of the inflammatory response.
- Oligodendrocyte Precursor Cells (OPCs): These agents are present in the parenchyma but passive in the current version of the BBBABM (Chodobski, Zink et al. 2011, Galea 2021, Nguyen, Pham et al. 2026).

#### 2.1.4 Extracellular and Soluble Mediators

The agents in the BBBABM existing in an environment that consists of overlapping fields of extracellular mediators. The variability of these fields is what produces the local heterogeneity that drives differential behavioral trajectories amongst the cellular agents, again mimicking how actual multicellular systems work. The BBBABM includes 18 soluble mediators/entities represented as model variables involved in BBB function. These entities, their role in the BBBABM and justifying references are seen in Table 1A-F.

**Table 1A-F:**
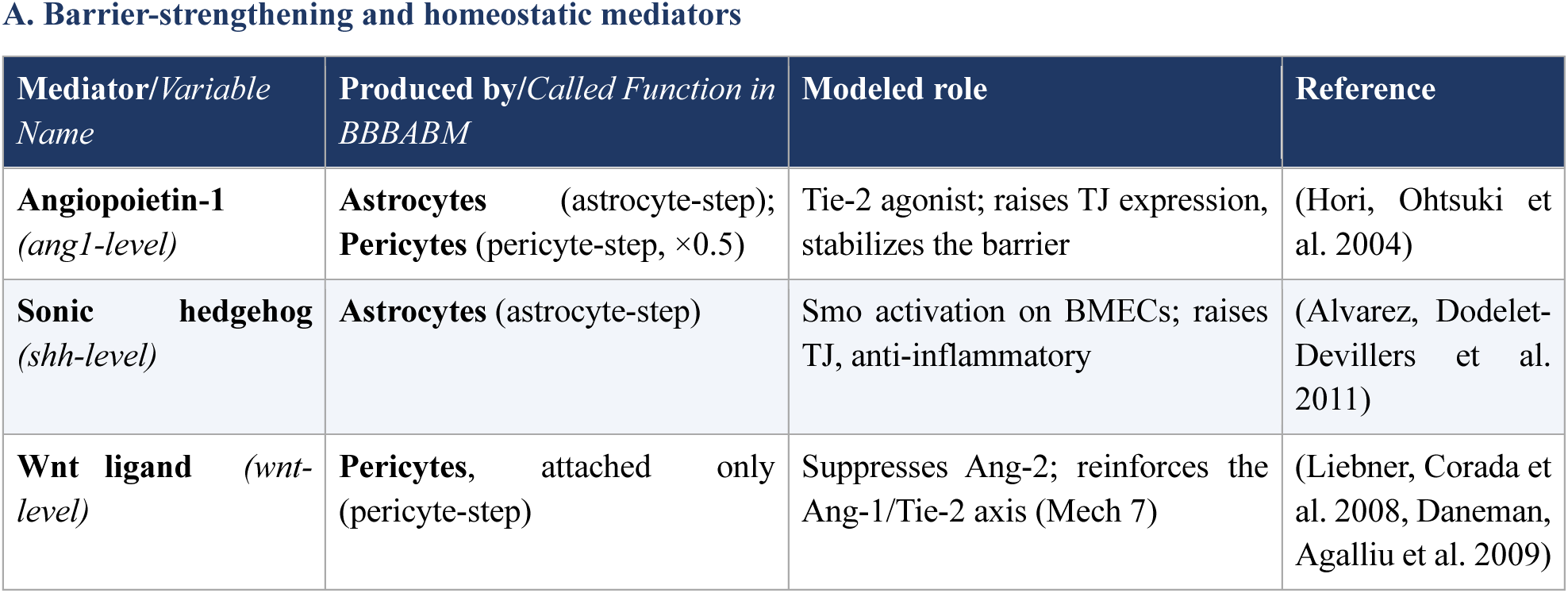
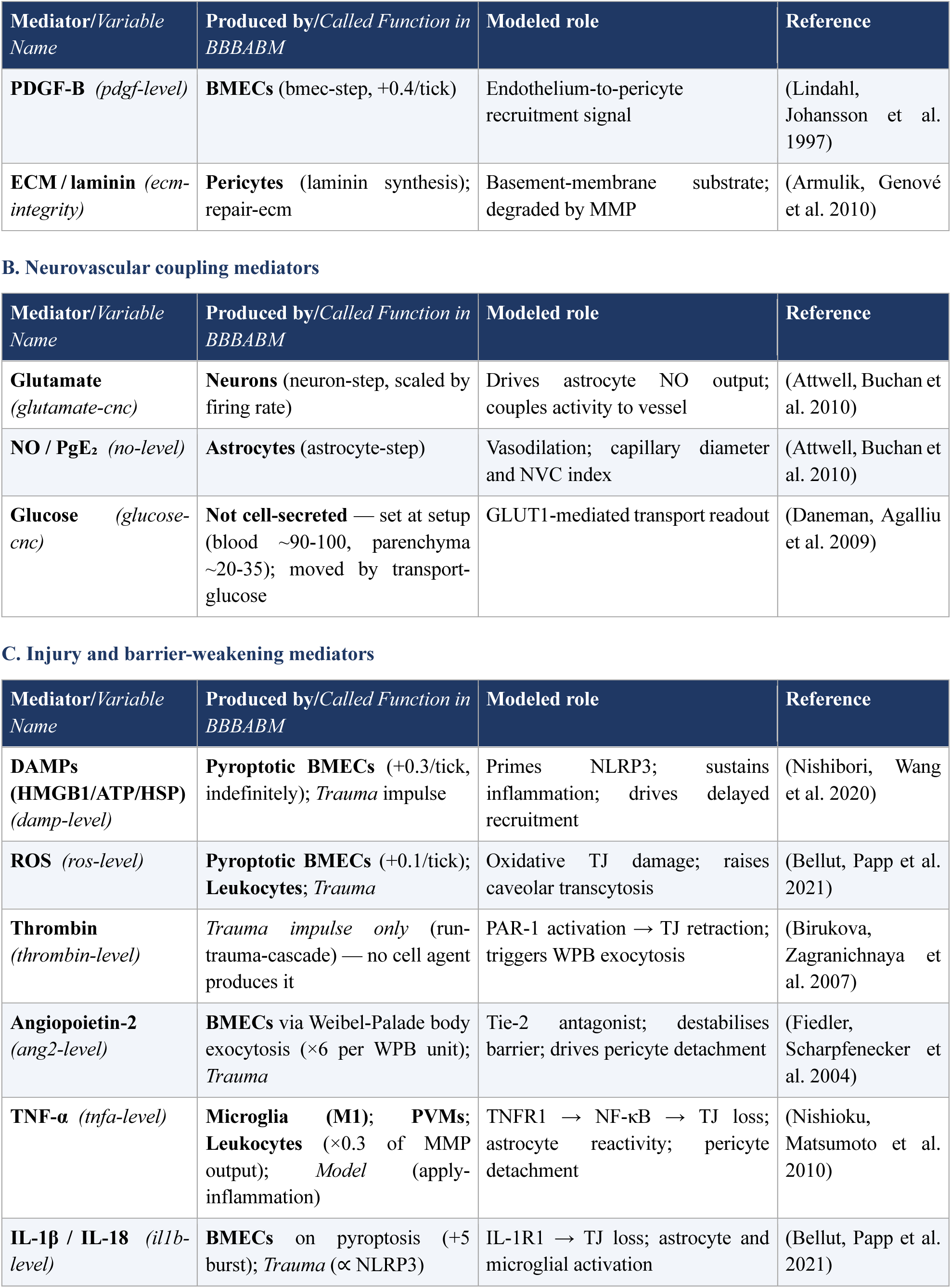
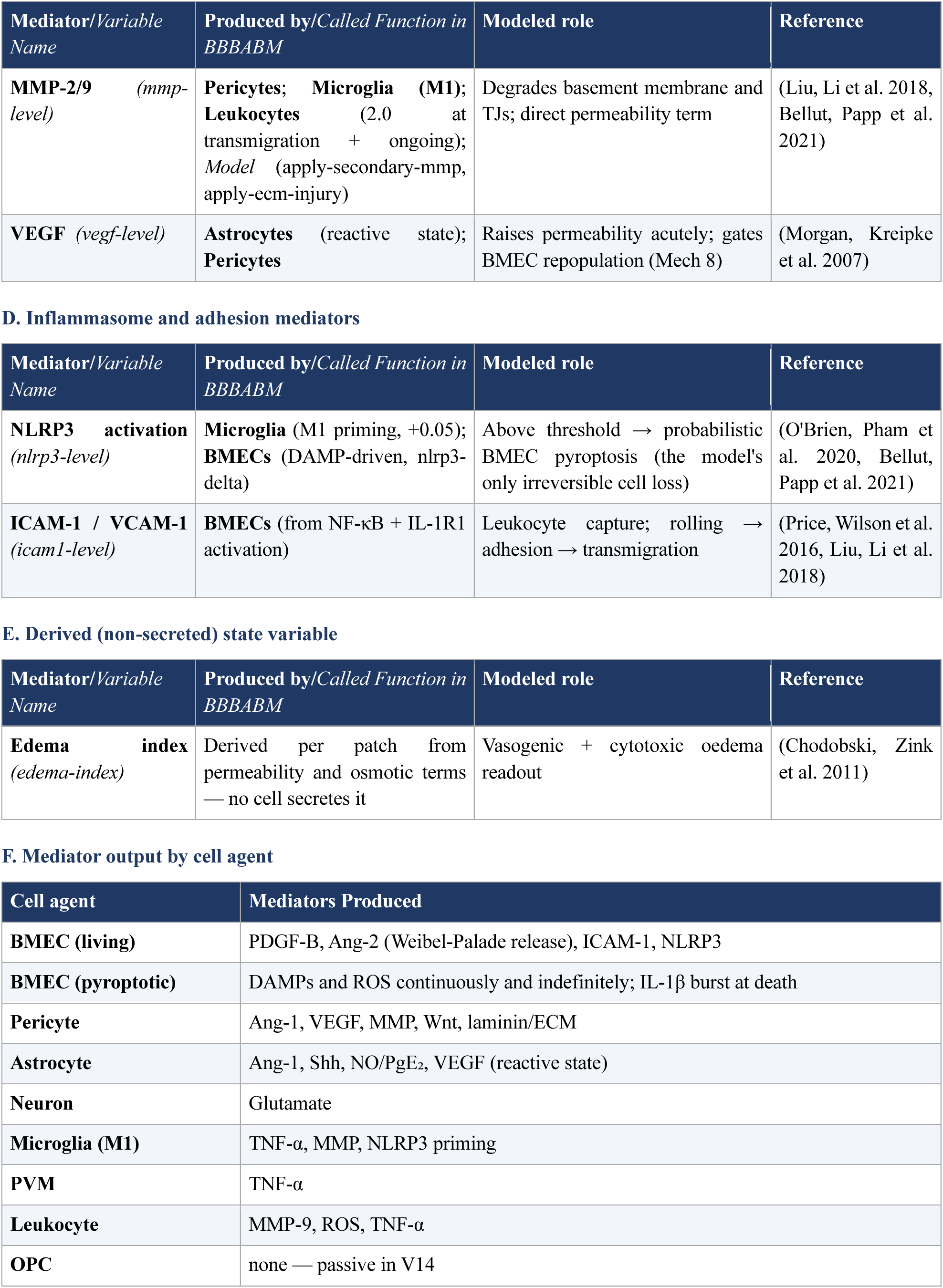
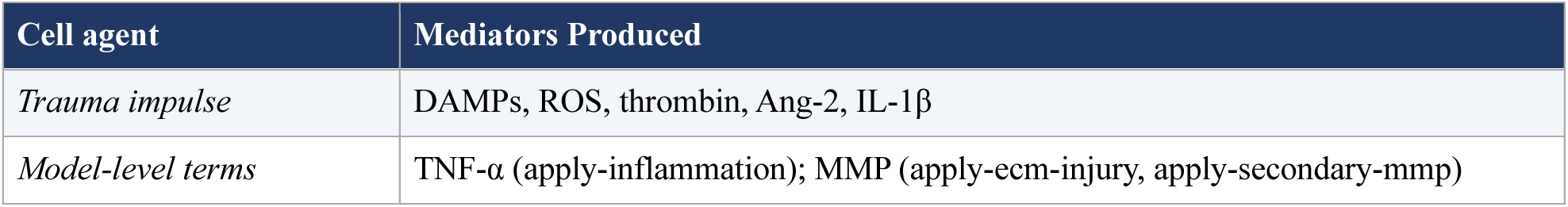
List of mediators and extracellular entities and their associated implementation labels in the BBBABM. *Notation:* **BMEC** = brain microvascular endothelial cell; **PVM** = perivascular macrophage; *Trauma* = the exogenous run-trauma-cascade function (not a cell agent); *Model* = a system-level term (apply-inflammation, apply-injury, apply-secondary-mmp) applied to the overall simulation model rather than an individual cell/agent.

#### 2.1.5 Mechanisms of BBB Injury

The following sections describe the various mechanisms by which the BBB can be disrupted. Each of these mechanisms are incorporated into the BBBABM and represent the processes that are compensated for in the restorative functions described in Section 3.1.6. In the simulation, the activation of these processes, the “trauma impulse”, are tied to a term called “*Trauma Severity*” which represents the extent of insult applied in the simulation: the parameter sweep used in the validation simulations varies this term to demonstrate progressively more severe insults.

##### 2.1.5.1 Mechanical Disruption of the Endothelial Monolayer

Mechanical force (contusion, shear, blast wave, etc.) directly fragments the endothelial cytoskeleton, tears intercellular junctions, and causes focal necrosis of BMECs. This produces an immediate paracellular gap that is independent of any subsequent signaling cascade — a point worth stating explicitly, since it means that no amount of anti-inflammatory intervention, however well-targeted, can address this means of BBB disruption. The molecular consequences of mechanic force injury are:

- **Claudin-5 loss:** Within 30 minutes of TBI, claudin-5 protein is lost from TJ complexes in peri-injury vessels, measurable by Western blot and immunofluorescence. This precedes any detectable neuroinflammatory signal (Chodobski, Zink et al. 2011, Price, Wilson et al. 2016).
- **Occludin phosphorylation:** Mechanical stress activates Src kinase, which phosphorylates occludin at Tyr398/402 and disrupts its interaction with ZO-1, causing TJ strand disassembly (Chodobski, Zink et al. 2011, Price, Wilson et al. 2016, Gopal, Kaur et al. 2025).
- **Cytoskeletal contraction:** Rho-ROCK signaling downstream of mechanical stress contracts the actomyosin cytoskeleton, pulling ZO-1/2 scaffolds away from the plasma membrane and widening paracellular gaps (Chodobski, Zink et al. 2011, Price, Wilson et al. 2016, Gopal, Kaur et al. 2025).

In the BBBABM this is implemented as a direct, signaling-independent decrement of TJ-expression at trauma-site BMECs scaled by the trauma-severity parameter, applied at the moment of injury in the simulation. The signaling-independence of this term is its most important property: it is the one component of the lesion that no downstream intervention in the model can prevent and therefore places a constraint on the completeness of any post-injury therapy; however, it does not necessarily preclude the potential of a prophylactic therapy to mitigate these processes.

##### 2.1.5.2 Ischemia-Reperfusion and Reactive Oxygen Species (ROS)

ROS arise from mitochondrial dysfunction during ischemia and, consequently, from the oxidative burst accompanying reperfusion. ROS act on the BBB through at least two distinguishable routes: direct oxidation of TJ proteins, and activation of latent MMP zymogens (Chodobski, Zink et al. 2011, Price, Wilson et al. 2016, Cash and Theus 2020). The BBBABM represents ROS as a rapidly diffusing, rapidly decaying patch field that both decrements TJ expression directly and increases caveolar transcytosis — the latter representing the transcellular component of barrier failure that is frequently neglected in accounts focused exclusively on paracellular leak.

We note, because it becomes important in Section 3.1.5.7 “Pyroptosis,” that in the BBBABM the dominant late source of ROS is not from the acute injury burst but rather is produced by the population of pyroptotic BMECs, which continue to release ROS indefinitely until cleared.

##### 2.1.5.3 Coagulation Cascade Activation: Thrombin and PAR-1

Disruption of the BMEC layer exposes subendothelial tissue factor and activates the coagulation cascade, generating thrombin at the injury site. Thrombin acts on endothelial protease-activated receptor 1 (PAR-1), triggering RhoA-mediated cytoskeletal contraction and TJ retraction (Chodobski, Zink et al. 2011, Price, Wilson et al. 2016, Cash and Theus 2020). Thrombin additionally activates microglia directly (Möller, Hanisch et al. 2000) and has been characterized across ischemic, hemorrhagic, and traumatic brain injury as having both deleterious and protective components depending on concentration (Xi, Reiser et al. 2003).

The BBBABM currently does not include the full coagulation cascade and represents thrombin as an exogenous mediator released by the trauma impulse rather than by any specific cellular agent.

##### 2.1.5.4 Angiopoietin-2 as a Trauma-Specific Barrier Destabilizer

Angiopoietin-2 (Ang-2) is stored preformed in endothelial Weibel-Palade bodies alongside von Willebrand factor and is released within minutes of stimulation by thrombin (Fiedler, Scharpfenecker et al. 2004). The fact that Ang-2 is already constitutively produced and stored for rapid release is what makes Ang-2 a trauma-specific rather than a generically inflammatory mediator and is therefore the reason it is treated separately from the cytokine cascade.

Released Ang-2 competes with Angiopoietin-1 at the endothelial Tie-2 receptor. Because Ang-1/Tie-2 signaling is the principal paracrine axis by which pericytes and astrocytes maintain junctional gene expression, Ang-2 release converts a pericyte-supported endothelium into an unsupported one without any change in pericyte behavior at all. The model implements this as competitive occupancy at Tie-2, with the resulting Ang-2/Ang-1 ratio serving as one of the readouts of barrier destabilization (Chodobski, Zink et al. 2011, Price, Wilson et al. 2016, Cash and Theus 2020).

##### 2.1.5.5 The Role of TGF-β

TGF-β is released in large quantities from aggregating platelets after vascular injury and is additionally synthesized by astrocytes (all three isoforms) and, to a lesser but still substantial extent, by microglia (Chodobski, Zink et al. 2011, Price, Wilson et al. 2016, Cash and Theus 2020). Astrocyte- and pericyte-derived TGF-β is required to enhance and maintain endothelial barrier properties, and that targeted disruption of Smad4 in brain endothelial cells — removing the capacity to respond to TGF-β — itself causes BBB breakdown by destabilizing the N-cadherin-dependent interaction between endothelium and pericytes (Li, Lan et al. 2011). In the BBBABM TGF-β serves as a pericyte-derived barrier-supporting term. There are additional potential roles for TGF-β that involve both supporting and disturbing the BBB, but these involve engaging more remote aspects of the inflammatory response (both pro- and anti-) and are not currently implemented in the BBBAB.

##### 2.1.5.6 Damage-Associated Molecular Patterns and the NLRP3 Inflammasome

Necrotic BMECs and surrounding parenchymal cells release damage-associated molecular patterns (DAMPs) into the interstitium within minutes of injury. The principal DAMPs relevant to BBB disruption are listed in Table 2.

**Table 2:** DAMPs associated with BBB dysfunction.

| DAMP | Origin | Pattern recognition receptor | Effect on BBB |
| --- | --- | --- | --- |
| **HMGB1** | Necrotic BMECs and neurons | TLR2, TLR4, RAGE | NF-κB activation → ICAM-1↑, TJ↓ |
| **ATP (extracellular)** | Necrotic cells (cytoplasmic release) | P2X7R on microglia, BMECs | NLRP3 inflammasome activation, K<sup>+</sup> efflux |
| **Heat-shock proteins (HSP60, HSP70)** | Stressed/necrotic cells | TLR2, TLR4 | Microglial TLR4 signaling → TNF-α, IL-6 |
| **Mitochondrial DNA** | Disrupted mitochondria | TLR9, cGAS-STING | Interferon-I response; NF-κB → cytokine storm |
| **S100B** | Astrocytes (necrotic/reactive) | RAGE on BMECs | RAGE → MAPK → TJ phosphorylation and loss |

HMGB1 is the representative member of this class and the one for which the barrier consequences are best characterized: released from injured neurons and glia, extracellular HMGB1 disrupts the BBB through endothelial contraction, microglial activation, and astrocytic endfoot swelling, producing edema as a composite consequence (Nishibori, Wang et al. 2020). In the current version of the BBBABM this heterogeneous group of DAMPs are consolidated into a single “*DAMP*” variable.

Extracellular ATP binding to P2X7R on BMECs and microglia triggers potassium (K⁺) efflux, providing the second signal for NLRP3 inflammasome assembly. The canonical pathway proceeds through four sequential steps:

1. **Signal 1 (priming):** HMGB1-TLR4-NF-κB transcription upregulates NLRP3 and pro-IL-1β protein in BMECs and microglial cells within 1–2 hours.
2. **Signal 2 (activation):** ATP-P2X7R-mediated K⁺ efflux, ROS, or lysosomal damage drives NLRP3 oligomerization, ASC speck formation, and caspase-1 recruitment.
3. **Caspase-1 cleavage:** Active caspase-1 cleaves pro-IL-1β → mature IL-1β, and pro-IL-18 → IL-18.
4. **Gasdermin D (GSDMD) pore formation:** Caspase-1 also cleaves GSDMD, inserting pores in the BMEC plasma membrane. At sub-lytic concentrations, GSDMD pores act as IL-1β release channels; at higher concentrations, they drive pyroptotic cell death.

A key point is that the endothelium itself, distinct from signaling in the microglial, executes this pathway: BMECs express NLRP3, upregulate it under injury, and undergo caspase-1- and gasdermin-D-dependent pyroptosis, with NLRP3 inhibition preserving ZO-1 staining and reducing albumin extravasation in vivo (Bellut, Papp et al. 2021). The same work demonstrates that endothelial NLRP3 activation drives MMP-9 secretion, which supplies the mechanistic link between inflammasome activity and the protease arm treated in Section 3.1.5.9.

##### 2.1.5.7 Pyroptosis and Irreversible Endothelial Cell Loss

Pyroptosis is placed into its own subsection, rather than treating it as the terminal step of Section 3.1.5.6. This is because BMEC pyroptosis is a categorially different type of event that produces long term effects affecting the population level properties of the BBBABM that to a great degree bridge the time scales between acute signaling events and those related to quasi-steady functions based on cell-state.

The initiating steps of pyroptosis come from NLRP activation (Section 3.1.5.6); as implemented in the BBBABM when local NLRP3 activation exceeds a set threshold, a BMEC undergoes pyroptosis with a per-tick probability scaling with the degree of threshold exceedance and with the pyroptosis-sensitivity parameter (parameters that are defined at initialization of the model). A pyroptotic BMEC ceases all barrier function but it is not removed from the simulation. It persists as a permanent gap in the BBB, without the capability to regenerate its TJs but rather continues to release DAMPs and ROS onto its patch every step for the duration of the simulation run. Three consequences follow that are visible in the model output.

First, the system-level *tight-junction-expression* barrier measure is bounded above by cell survival. Because the reported mean is taken across all 33 endothelial positions including the dead cells the final value is quantized in units of 1/33: it equals (33 − *n*_pyroptotic_)/33. In parameter sweeps this quantization is exact — no run terminates at a non-quantized value, which is to say that no surviving cell ever has partial junctional integrity.

Second, there is a sharp severity threshold rather than a graded dose-response. Across trauma-severity values from 0.2 to 1.0 with ten replicates each, no endothelial cells die below approximately 0.6, and the barrier recovers completely. Above that threshold, cell loss rises steeply — roughly 1 cell at *trauma severity* 0.6, 9 at 0.7, and 12–15 at 0.9–1.0 — and the barrier plateaus permanently below baseline. The threshold region is also the region of greatest run-to-run variance, which is what one would expect of a stochastic threshold crossing and which we regard as a modest internal validation that the mechanism is behaving as a threshold rather than as a rescaled continuum.

Third, the observed persistent inflammatory signature in the BBBABM simulations is a corpse signature. Total DAMP does not decay to zero in any run where cells died, because dead cells never stop emitting. The residual DAMP plateau is therefore a direct readout of how many cells were lost, and — as Section 3.3 develops — it is also what supplies the delayed inflammatory drive underlying the second permeability peak.

We draw attention to this cluster of findings because it inverts a natural reading of the acute-injury literature. The intuition that a more severely injured barrier is one whose junctions are more severely damaged turns out, in this model, to be wrong in an interesting way: junctional damage is fully reversible in every surviving cell at every severity tested. What is not reversible is the loss of the cell itself, until the process of endothelial regeneration occurs (See Section 3.1.6.8 below).

##### 2.1.5.8 Blood-Borne Factors: Fibrinogen and Albumin as Independent Amplifiers

Once the BBB loses its integrity and becomes permeable, certain plasma proteins entering the parenchyma can become active signaling rather than passive markers of leak. These include:

- **Fibrinogen:** Beyond its coagulation role, fibrinogen deposited in the parenchyma activates microglia through CD11b/CD18 and, independent of microglial action, inhibits neurite outgrowth by acting as a ligand for αVβ3 integrin and transactivating the neuronal EGF receptor (Schachtrup, Ryu et al. 2010) — a direct mechanistic link between barrier leak and impaired neuronal repair that does not require any inflammatory intermediate. Fibrinogen additionally serves as a carrier for latent TGF-β and, upon fibrin conversion and TGF-β activation, promotes astroglial scar formation (Schachtrup, Ryu et al. 2010).
- **Albumin:** Albumin increases microglial intracellular calcium and drives proliferation, and in both microglia and astrocytes activates MAPK signaling and IL-1β synthesis (Ralay Ranaivo and Wainwright 2010). Its more consequential action is on astrocytes: albumin binds TGF-β receptor II directly and activates Smad signaling, and this albumin-dependent TGF-β pathway has been mechanistically implicated in post-traumatic cortical epileptogenesis (Ivens, Kaufer et al. 2007, Cacheaux, Ivens et al. 2009) — a chronic neurological sequela whose origin traces back to an acute failure of paracellular exclusion rather than to any neuronal injury mechanism per se.

We note these actions specifically because they illustrate that BBB breakdown is not merely a permeability defect to be corrected, but an active source of novel, parenchyma-directed signaling whose consequences can be entirely independent of — and can outlast — the inflammatory cascade with which it is usually grouped. These mechanisms have potential implications for chronic recurrent BBB dysfunction as might be seen in Alzheimer’s Disease, Multiple Sclerosis or CTE. At the moment, neither fibrinogen nor albumin are explicitly represented in the BBBABM (given the time scale of current interest); we note these factors as they represent the next targets for model enhancement.

##### 2.1.5.9 Leukocyte Recruitment and Transmigration

Endothelial activation upregulates ICAM-1 and VCAM-1, converting the luminal surface into an adhesive substrate. Circulating neutrophils, the most abundant leukocyte in circulation and the first transmigrating population at sites of injury, roll, adhere, and cross into the parenchyma, where they release MMP-9 and reactive oxygen species (Liu, Li et al. 2018). ICAM-1 and ICAM-2 mediate the attachment step, and MMPs are required for penetration of the parenchymal basement membrane (Price, Wilson et al. 2016).

Neutrophil-derived MMP-9 is the principal effector of secondary, protease-mediated barrier damage: it degrades type IV collagen in the basal lamina and cleaves tight-junction proteins directly. The distinction between this protease-mediated damage and the primary mechanical lesion of Section 3.1.5.1 matters therapeutically, since the former is in principle interruptible on a clinically relevant timescale and the latter is not.

In the BBBABM the leukocyte agents implement the full rolling-adhesion-transmigration sequence and deposit MMP at the endothelium on crossing. It should be noted that in the current BBBABM BMECs also constitutively produce MMP.

##### 2.1.5.10 Cerebral Edema: Vasogenic and Cytotoxic Mechanisms

Edema in the BBBABM is not a mediator but a derived value resulting from the mechanisms implemented in the model and therefore represents a model output that is recorded. Vasogenic edema follows from paracellular and transcellular barrier failure permitting oncotically active plasma protein into the interstitium; cytotoxic edema follows from failure of cellular ion homeostasis, principally through AQP4 dysregulation and Na⁺/K⁺-ATPase failure, and occurs with an intact barrier (Chodobski, Zink et al. 2011, Price, Wilson et al. 2016, Cash and Theus 2020).

#### 2.1.6 Mechanisms of Recovery: Eight Parallel, Redundant Processes

The brain endothelium possesses several intrinsic and paracrine repair mechanisms that act on distinct timescales — from minutes (local junctional resealing) to days and weeks (cellular replacement) — and that are mechanistically distinct from the pathways that produced the injury in the first instance. The BBBABM implements eight such mechanisms. We present them here in ascending order of characteristic timescale, and we note that this ordering is also approximately the order in which they become rate-limiting: a barrier that fails to reseal in the first hour fails for reasons involving Mechanisms 1 and 2, whereas a barrier still deficient at two weeks fails for reasons involving Mechanism 8 alone.

##### 2.1.6.1 Mechanism 1 — Rho-Flare Local Junctional Resealing (minutes)

At the single-junction level, transient localized bursts of RhoA activity — termed “Rho flares” — are triggered at the exact site of a tight-junction break by mechanosensitive calcium influx and p115RhoGEF recruitment. We draw particular attention to the fact that this is the same RhoA pathway implicated in the pathological, sustained contraction described in Section 3.1.5.1; the distinguishing feature is not the molecule but its spatial and temporal signature. Unlike the diffuse, sustained RhoA/ROCK activation that drives pathological endothelial contraction during the injury phase, Rho flares are spatially confined, resolve within minutes, and produce localized actomyosin accumulation that re-concentrates claudin-5, occludin, and ZO-1 at the breach site (Stephenson, Higashi et al. 2019, Chumki, van den Goor et al. 2022, Varadarajan, Chumki et al. 2022).

This is the fastest-acting repair mechanism and, notably, operates as a continuous, low-level surveillance-and-patch process even in uninjured endothelia — it is not a repair mechanism switched on by trauma so much as a constitutive maintenance process whose local amplitude scales with local damage. The model implements it as a damage-triggered flare with probabilistic initiation, fixed duration, and a per-tick junctional increment, active only in cells whose tight-junction expression fell on the preceding tick.

##### 2.1.6.2 Mechanism 2 — cAMP/Rac1 Counter-Regulatory Signaling (tens of minutes)

At the whole-cell level, recovery from TJ-disrupting stimuli such as thrombin requires active counter-signaling rather than simple decay of the injurious stimulus — a distinction we consider essential, since a model that represents recovery only as the passive decay of an injury signal will systematically overestimate how long barrier disruption should persist. Endogenous cyclic AMP signaling activates Rac1, a GTPase antagonizing RhoA-driven contraction and driving reassembly of the cortical actin ring that anchors tight junctions. This Rac1-dependent barrier restoration begins within minutes of thrombin exposure and is the dominant mechanism by which endothelial monolayers recover transendothelial electrical resistance even while thrombin remains present (Birukova et al. 2007; Aslam et al. 2014).

It is worth being precise about what this mechanism does in the model, because the natural shorthand overstates it. cAMP accumulates in proportion to PAR-1 activation and then scales the PAR-1 retraction term by (1 − cAMP). The effect is therefore attenuation of one specific injury term, bounded below by zero: at saturation the thrombin contribution is fully suppressed, but the term never reverses sign, and no other injury input — TNF-α, MMP, IL-1β, ROS — is damped by it at all. Mechanism 2 does not reverse retraction; it caps how far thrombin alone can drive it. That the same signal which opens the junction also builds the brake that limits it is the structural reason a single thrombin insult is self-limited rather than progressive, and — by direct implication — why loss of this specific counter-regulatory capacity, rather than simply “more injury,” is a plausible route by which an otherwise survivable insult becomes a progressive one.

##### 2.1.6.3 Mechanism 3 — The Biphasic Permeability Curve (hours to days)

*In vivo*, BBB permeability after TBI follows a well-documented biphasic pattern rather than a single monotonic recovery: an early permeability peak at approximately 4–6 hours post-injury, a partial closure, and a second peak at approximately 2–3 days (Başkaya et al. 1997). TJ protein levels track this curve, with blast-exposure studies reporting substantial normalization by roughly 3 days and stability thereafter (Kawoos et al. 2021).

During the development of the BBBABM the intent was to have this biphasic behavior emerge from the interaction of fast-acting resealing mechanisms (Mechanism 2) with slower leukocyte and protease dynamics. However, this did not work, and the reasons it did not work are more instructive than the position we originally took.

The emergent approach failed for a structural reason rather than a parametric one. The second peak requires a driver arriving after the first has resolved; the leukocyte module, however, was gated on endothelial ICAM-1, which decays with the acute inflammatory burst and is functionally absent within roughly fifteen hours. Recruited leukocytes died within 150 ticks of arrival. The entire cellular arm therefore completed its work approximately twenty-fold earlier than the phenomenon it was meant to produce, and no adjustment of recruitment amplitude could move an event whose *timing* was determined by the decay constant of a fast-decaying field. Successive attempts to repair this by amplifying the driving signal revealed a second and more general problem: the path from injury signal to permeability response ran through eight sequential multiplicative stages — driver, recruitment, spawning, adhesion, crossing, deposition site, lateral diffusion, and permeability coupling — such that the end-to-end effect was the product of eight small factors and was repeatedly negligible.

The current BBBABM therefore implements Mechanism 3 through an approach that more directly encodes the time delays needed for the early resolution and delayed permeability failure reported in the literature. For this we utilize the pyroptotic state of the BMECs, where there is an intracellular buildup of DAMP, whose magnitude scales of *Trauma Severity*, that is then released in a delayed fashion as pyroptosis progresses resulting in a delayed recruitment drive within a 12-to-48-hour window, which in turn drives a direct MMP-9 burst at the BMEC layer. The result is a genuine second peak: at trauma-severity 0.5–0.6 the model produces an acute peak near 30 minutes, a trough at approximately one day, a second peak at approximately two days, and resolution by day four, with second-peak magnitude scaling monotonically with severity.

One further observation merits mention, because it was not anticipated. The clean biphasic curve appears at moderate severity (0.5–0.6) and degrades at high severity — not because the second wave weakens, but because the trough disappears. Above the pyroptosis threshold, cell loss holds mean permeability elevated, so the second peak reads as a step on a plateau rather than as a separate excursion. The biphasic signature is thus a property of injuries survivable at the cellular level, which is a prediction the model makes and which we have not seen stated in the experimental literature.

##### 2.1.6.4 Mechanism 4 — Claudin-5 Transcriptional Upregulation and Overshoot (days to weeks)

Following the acute resealing phase, claudin-5 transcription is upregulated above baseline: experimental TBI models show claudin-5 expression increasing 1–2 weeks after injury and remaining elevated for as long as 4–8 weeks (Price, Wilson et al. 2016). The BBBABM implements a transcription-rate variable that rises toward a ceiling while trauma signals persist and relaxes slowly thereafter, contributing an additive term to junctional synthesis.

##### 2.1.6.5 Mechanism 5 — MMP-9 Resolution via the CypA Pathway (days)

Restoration of TJ protein stability is determined by the decline of active MMP-9, which is otherwise sustained by a cyclophilin A (CypA)-dependent pathway in the injured neurovascular unit. Pharmacological inhibition of CypA attenuates MMP-9 activity and measurably accelerates spontaneous BBB repair after TBI in mouse models, demonstrating that the rate of MMP-9 resolution — not merely its peak level — sets the pace of junctional restabilization (Main, Villapol et al. 2018). This places MMP-9 clearance, alongside RhoA inactivation, as a second rate-limiting node in TJ recovery, whose kinetics, not merely its presence or absence, constitutes the actual target of intervention. The model exposes this as a tunable clearance rate representing genotype-dependent repair speed, apoE3-like at high values and apoE4-like at low.

##### 2.1.6.6 Mechanism 6 — Pericyte Repopulation and Paracrine Restabilization (days)

Pericyte coverage is acutely reduced after TBI — up to 40% of pericytes lose contact with the basement membrane within the first hours — and its restoration is necessary for full BBB stabilization, since pericytes are the dominant paracrine source of Ang-1 and TGF-β acting on the endothelium. Pericyte-derived Ang-1 forms a multimeric complex activating endothelial Tie-2 and directly inducing occludin transcription, while pericyte TGF-β independently upregulates BBB-specific gene expression (Hori, Ohtsuki et al. 2004, Dohgu, Takata et al. 2005, Armulik, Genové et al. 2010).

##### 2.1.6.7 Mechanism 7 — Wnt/β-catenin-Driven Junctional Gene Re-Expression (ongoing)

Independent of the acute repair mechanisms above, sustained canonical Wnt/β-catenin signaling — delivered to brain endothelium by perivascular pericytes and CNS neural cells via Wnt7a/7b ligands acting on endothelial Frizzled/LRP5/6 receptors — drives transcriptional induction of BBB-specific genes, including downregulation of the Tie-2 antagonist Ang-2 and upregulation of Ang-1 (Liebner, Corada et al. 2008, Daneman, Agalliu et al. 2009). We emphasize that this pathway functions less as a discrete repair switch than as a continuously active maintenance signal: its restoration, following any disruption of pericyte or neuronal Wnt ligand supply, is required for durable rather than transient barrier competence. In the model it is implemented as continuous secretion by attached pericytes with a threshold-gated suppression of Ang-2, which makes Mechanism 7 dependent on Mechanism 6 — a coupling we consider biologically correct and computationally consequential, since it means pericyte loss degrades the barrier through two distinct routes rather than one.

##### 2.1.6.8 Mechanism 8 — BMEC Repopulation (days to weeks)

This mechanism addresses the recovery from pyroptotic BMECs, whose persistence would otherwise preclude recovery to baseline for even trivial insults. Quiescent brain endothelium turns over extremely slowly — estimates of endothelial turnover time in normal tissues range from 47 to 23,000 days, against potential doubling times of 2.4 to 13 days for tumor endothelium (Hobson and Denekamp 1984), which corresponds to a replacement rate on the order of 0–1% per day. After injury, however, proliferation is induced on a days-to-weeks clock: VEGF and VEGFR2 rise after TBI, with BrdU incorporation confirming newly formed vessels within 48 hours (Morgan, Kreipke et al. 2007), peak peri-lesional proliferation at approximately 3 days, and peak vessel density at approximately 14 days.

The BBBABM implements this as a per-tick probability that a pyroptotic cell is replaced from a living neighbor, scaled by local VEGF, with the baseline rate expressed in the literature’s own units (percent of the layer per day) and converted internally. Replacement cells enter with immature, leaky junctions and must then mature through Mechanisms 1 through 7 — a detail we consider important rather than decorative, since it means cellular replacement does not immediately restore barrier function and produces a second, slower recovery phase lagging behind cell census.

Figure 2 summarizes the BMEC intracellular mechanisms of TJ injury and restitution.

**Figure 2:**
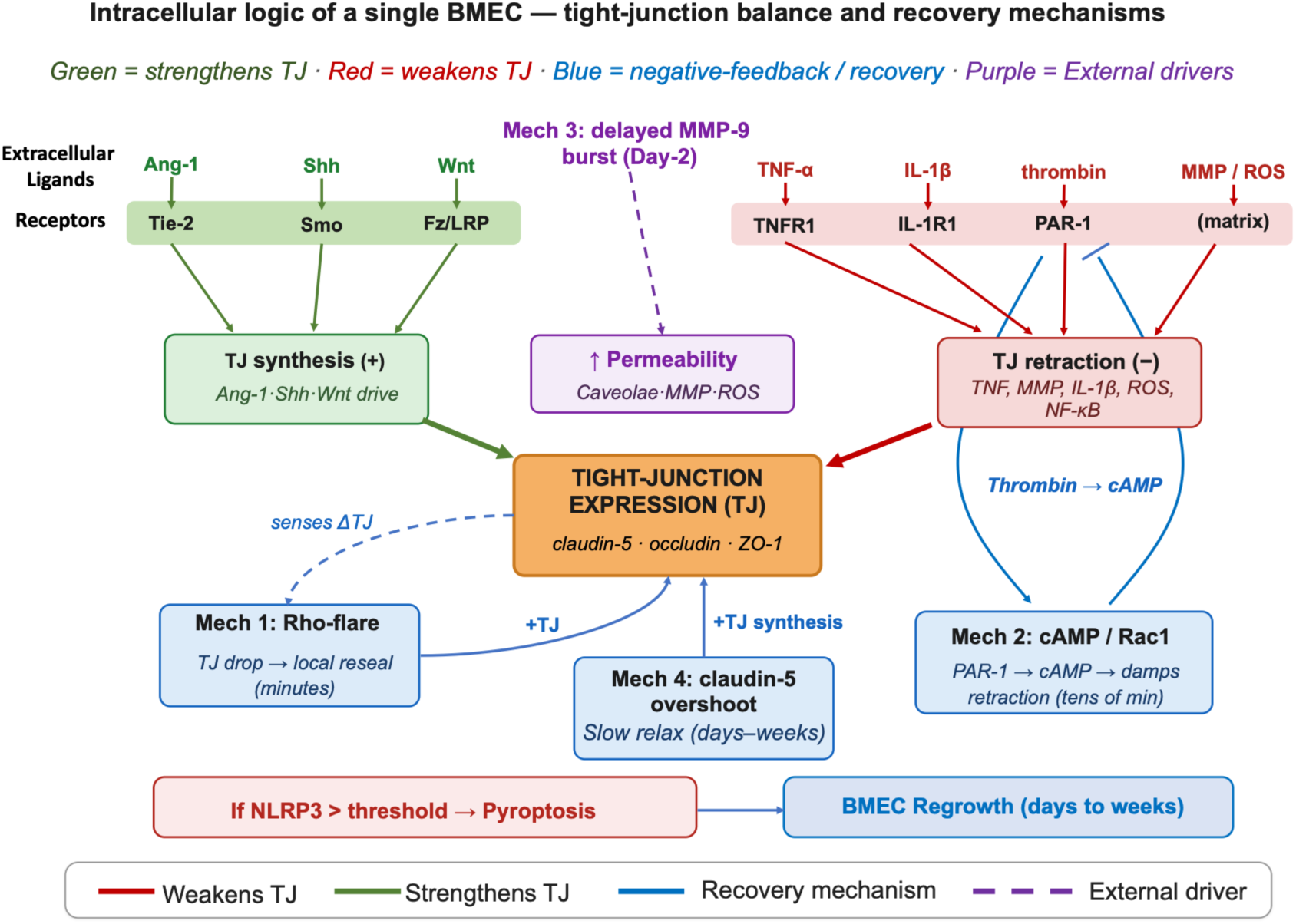
BMEC Intracellular interactions regarding TJ-expression.

### 2.2 Simulation results

As noted above the simulation experiments involved a parameter sweep of the *Trauma Severity* variable with application of the insult at T=0 and the duration of the simulation = 14 days. Figure 4A-B show the trajectories of TJ-expression across valued for *Trauma Severity* from 0.2 to 0.9 (dimensionless values). Each condition was run with 10 stochastic replicates; the solid line is the mean of those runs and the shaded area shows the distribution of the variable runs. Panel A shows the trajectories over the full 14 day run, while Panel B is a zoomed-in plot that emphasizes the period (∼ 33 hrs) where the primary dynamics. Both plots show that with *Trauma Severity* of 0.2 – 0.3 the degree of TJ disruption never maximizes within the context of the simulation and there is a rapid recovery to pre-insult levels. The degree of TJ disruption is small enough to show the dose-dependency of the disruption vs *Trauma Severity.* At *Trauma Severity* = 0.4 – 0.5 there is a period of maximal TJ disruption but this also recovers to baseline relatively quickly. At *Trauma Severity* = 0.6 – 0.9 there is more significant and prolonged disruption, and the lack of recovery before 2000 minutes is evidence that BMECs have undergone pyroptosis where baseline TJ expression cannot be recovered until BMEC replacement has occurred (Recovery Mechanism 8).

**Figure 3A-B.**
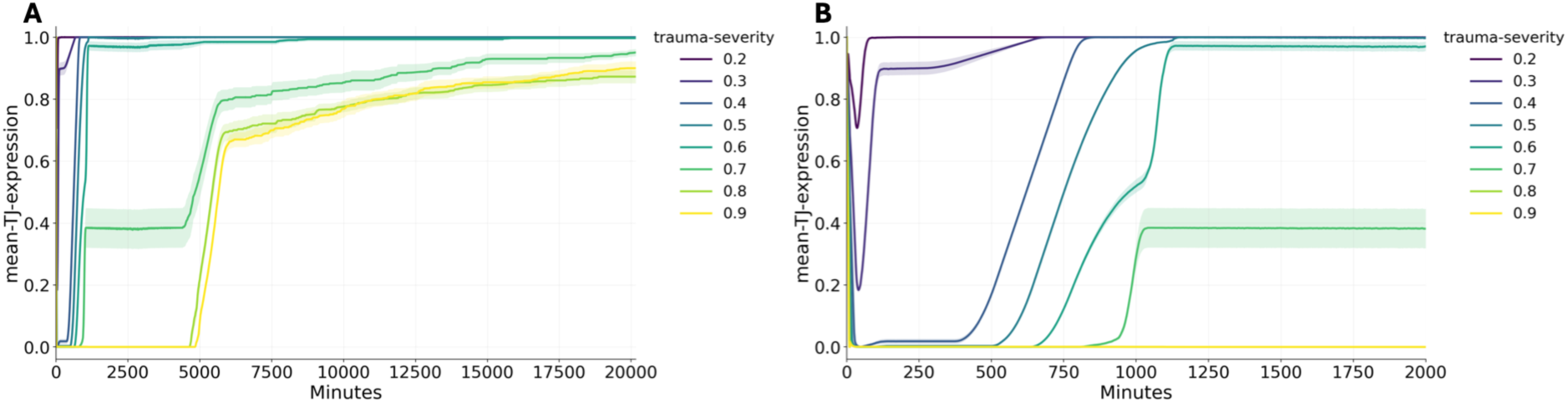
Mean TJ-expression trajectories at 14 days (Panel A) and ∼33 hours (Panel B)

Figure 4A-B shows the corresponding permeability dynamics, again with Panel A showing the 14-day run and Panel B showing the ∼33 hr run.

**Figure 4A-B.**
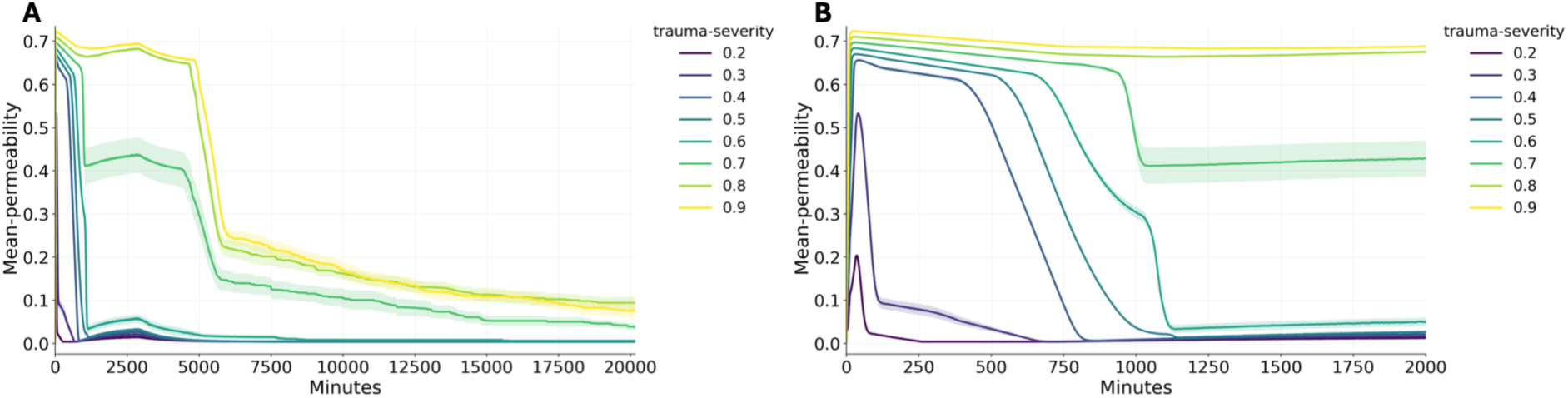
Mean permeability trajectories at 14 days (Panel A) and ∼33 hours (Panel B)

After trauma, permeability shows a sharp acute peak, then (at low-to-moderate severity) recovers into a trough, then rises again around day 2 in a second, smaller peak before resolving. This biphasic shape is the signature of Mechanism 3 and matches the two-phase barrier opening reported after TBI. The clearest biphasic curves appear at severities around 0.5-0.6. At higher severity the trough is shallow because dead cells keep the barrier open, so the second wave reads as a step rather than a separate hump.

To parse the BBB recovery dynamics Figure 5A-B compares the TJ-expression dynamics (Panel A) versus the number of pyroptotic BMECs (Panel B). As noted above, pyroptotic BMECs do not ever recover their TJ integrity; their contribution to increased permeability persists until they are replaced at the much longer time scale seen in Mechanism 8. As currently constructed, the BBBABM allows any BMECs that escape pyroptosis to eventually recover their TJ integrity. This is seen comparing Panel 5A with Panel 5B where the persistent TJ-expression deficit at high *Trauma Severity* is cell death, not failed junction repair; TJ-expression does not start to recover until the pyroptotic BMEC population starts to fall (evidence of BMEC regeneration).

**Figure 5A-B.**
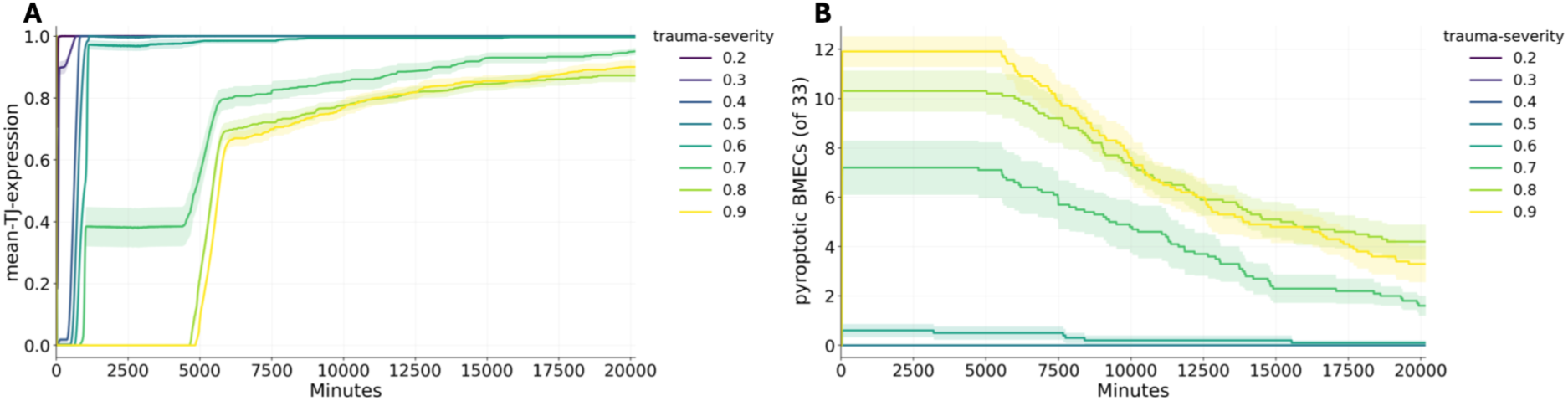
Panel A = TJ-expression over 14 days, Panel B = Pyroptotic BMECs over 14 days.

Neuroinflammation has been recognized as a significant contributing process to a host of neurological pathologies, and the endothelial activation that initiates BBB dysfunction involves inflammatory pathways. Figure 6 tracks the levels of inflammation-related metrics across the parameter sweep of *Trauma Severity*: TNF-*a*, leukocytes that have migrated into the CNS and MMP levels. These dynamics all show an initial inflammatory signal immediately following application of the perturbation, followed by a considerably larger spike consistent with the time delay associated with the process of leukocyte migration and their subsequent production of inflammatory mediators at around 2 days; it is this dynamic that generates the biphasic permeability pattern noted in Mechanism 3. What is notable is that even with high values of *Trauma Severity* there is no generation of a self-propagating neuroinflammatory loop; this is consistent with the putative pathophysiology of chronic neurological disorders that involves ongoing, recurring activation of the endothelium in the neurovascular unit.

**Figure 6A-C.**
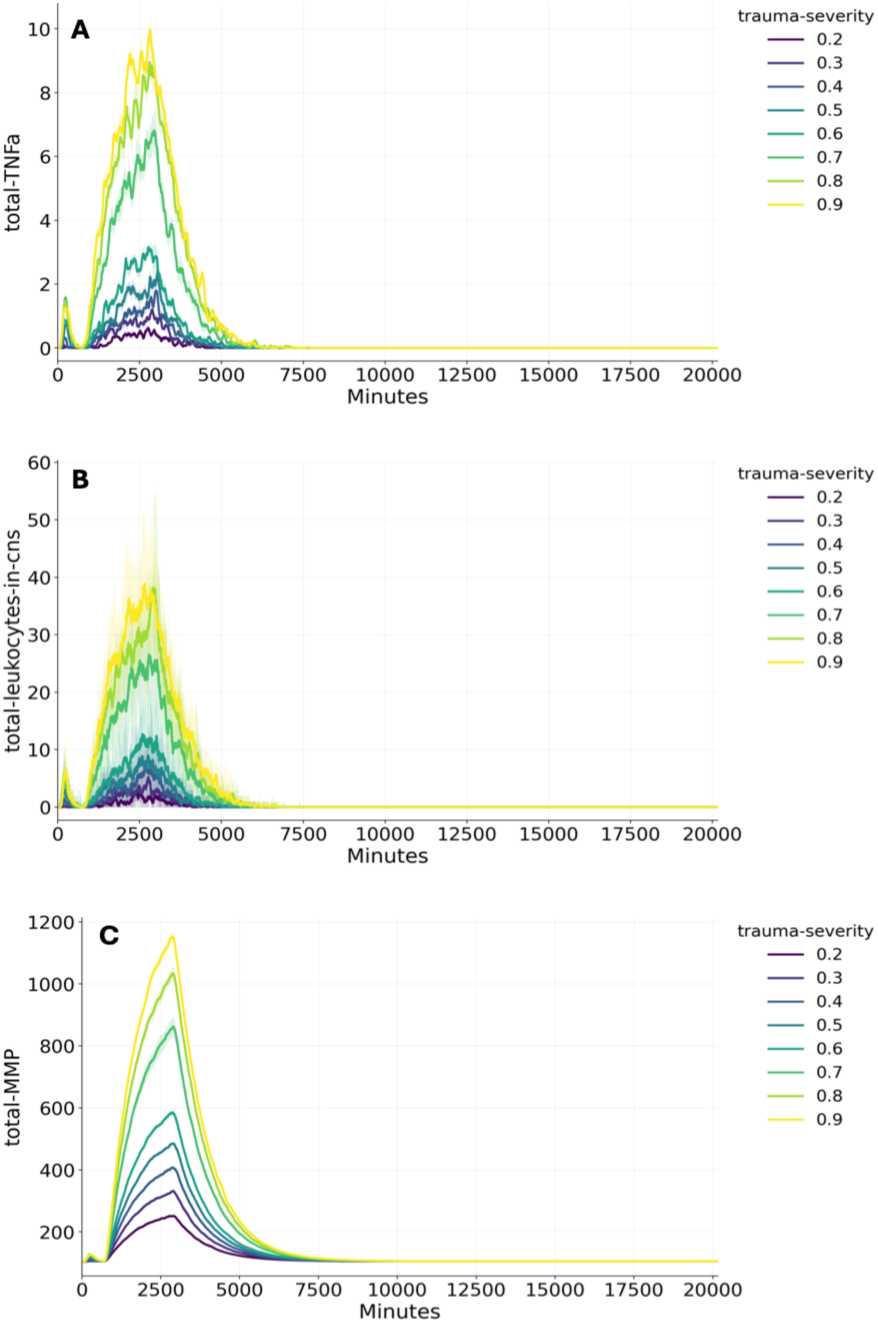
Trajectories of TNF-⍺ (Panel A), migrated leukocytes (Panel B) and MMP (Panel C)

## 3. Discussion

The BBBABM is, to our knowledge, the first computational model that incorporates the known cellular architecture of the neurovascular unit, the molecular mechanisms of BBB dysfunction and the mechanisms associated with BBB restitution (see Review from 2026 (Jafarabadi, Datta et al. 2026)). We believe that modeling disease always starts with modeling the state of health, and we recognize that the state of health requires representing processes and functions that maintain homeostasis and beneficial responsiveness to daily insults. While the initiating perturbation in the BBBABM is labeled “trauma” the modeled process of endothelial activation of the BBB is representative of what a real individual’s neurovascular unit would encounter while living in a variable and uncertain environment and reflects how that system would manifest its homeostatic mechanisms. From this point various disease states can emerge by altering the context and conditions applied to the BBABM, and therefore we believe that the BBBABM can readily be extended to simulation a host of neuro-pathologies known to involve dysfunction of the BBB: TBI (Shlosberg, Benifla et al. 2010, Chodobski, Zink et al. 2011, Hay, Johnson et al. 2015, Price, Wilson et al. 2016, Cash and Theus 2020, Sulhan, Lyon et al. 2020, van Vliet, Ndode-Ekane et al. 2020), CTE (Doherty, O’Keefe et al. 2016, Greene, Brennan et al. 2026), multiple sclerosis (Nishihara, Perriot et al. 2022, Zierfuss, Larochelle et al. 2024), Alzheimer’s Disease (Montagne, Zhao et al. 2017, Sweeney, Sagare et al. 2018) and stroke (Abdullahi, Tripathi et al. 2018, Yang, Hawkins et al. 2019). In addition, since the BBB is responsive to circulating inflammatory mediators (Elwood, Lim et al. 2017, Galea 2021, Huang, Hussain et al. 2021), including those seen in burn injuries (Yang, Ma et al. 2020, Xie, Hu et al. 2022, Chen, Zhang et al. 2023), the BBBABM can potentially be integrated into existing mult-organ/tissue ABMs of systemic inflammation (An 2008).

While the BBBABM currently focuses on the response of the neurovascular unit to a single perturbation, extensions in the near future will incorporate the secondary effects that are a consequence of acute permeability failure, specifically the inflammogenic properties of molecules such as albumin and fibrinogen, that can leak into the CNS parenchyma.

As a computational model, the BBBABM can be used to simulate the extended time scales (years to decades) involved in more chronic processes, providing a previously unavailable capability to mechanistically observed how these diseases evolve (see example regarding oncogenesis (An and Kulkarni 2015).

The modular and mechanistic nature of the BBBABM offers considerable opportunity in terms of integrating with other cutting-edge tools being used to study disease processes involving the BBB. With the increasing attention and interest in novel alternative methods (NAMS) (https://www.fda.gov/science-research/science-and-research-special-topics/new-approach-methodologies-nams , https://www.nih.gov/about-nih/who-we-are/nih-director/statements/statement-catalyzing-development-novel-alternatives-methods ), the structure of the BBBABM can be readily mapped to neuro-tissue BBB organoids (Bergmann, Lawler et al. 2018, Nzou, Wicks et al. 2018, Logan, Arzua et al. 2019) and BBB organ-on-a-chip systems (Kawakita, Mandal et al. 2022, Nasiri, Madadelahi et al. 2025, Vetter, Palagi et al. 2025). The BBBABM also has potential integration with physics-based models of TBI (Jafarabadi, Datta et al. 2026, Sasidharan, Tan et al. 2026). These models, in turn, can bridge the BBBABM with experimental “phantoms” of TBI to provide an unprecedented level of linkage across scales extending from molecular processes to anatomic (e.g. protective head-gear design).

Finally, as a mechanistic computational model, in addition to serving as a means of evaluating and refining a particular hypothesis structure (as was done during this project), the BBBABM can be used as a testing platform to discover new therapies and, perhaps more importantly, new combinations of therapies (Cockrell and An 2018), eventually into the context of a true mechanistic cellular-molecular based Digital Twin (An and Cockrell 2024, Cockrell, Vodovotz et al. 2024).

